# Rapid and memory-efficient analysis and quality control of large spatial transcriptomics datasets

**DOI:** 10.1101/2024.07.23.604776

**Authors:** Bence Kӧvér, Alessandra Vigilante

## Abstract

The 10x Visium spatial transcriptomics platform has been widely adopted due to its established analysis pipelines, robust community support, and manageable data output. However, technologies like 10x Visium have the limitation of being low-resolution, and recently spatial transcriptomics platforms with subcellular resolution have proliferated. Such high-resolution datasets pose significant computational challenges for data analysis, with regards to memory requirement and processing speed. Here, we introduce Pseudovisium, a Python-based framework designed to facilitate the rapid and memory-efficient analysis, quality control and interoperability of high-resolution spatial transcriptomics data. This is achieved by mimicking the structure of 10x Visium through hexagonal binning of transcripts. Analysis of 47 publicly available datasets concluded that Pseudovisium increased data processing speed and reduced dataset size by more than an order of magnitude. At the same time, it preserved key biological signatures, such as spatially variable genes, enriched gene sets, cell populations, and gene-gene correlations. The Pseudovisium framework allows accurate simulation of Visium experiments, facilitating comparisons between technologies and guiding experimental design. Specifically, we found high concordance between Pseudovisium (derived from Xenium or CosMx) and Visium data from consecutive tissue slices. We further demonstrate Pseudovisium’s utility by performing rapid quality control on large-scale datasets from Xenium, CosMx, and MERSCOPE platforms, identifying similar replicates, as well as potentially low-quality samples and probes. The common data format provided by Pseudovisium also enabled direct comparison of metrics across 6 spatial transcriptomics platforms and 59 datasets, revealing differences in transcript capture efficiency and quality. Lastly, Pseudovisium allows merging of datasets for joint analysis, as demonstrated by the identification of shared cell clusters and enriched gene sets in the mouse brain using data from multiple spatial platforms. By lowering the computational requirements and enhancing interoperability and reusability of spatial transcriptomics data, Pseudovisium democratizes analysis for wet-lab scientists and enables novel biological insights.

## Introduction

Spatial transcriptomics localises and quantifies RNA species in a tissue sample (1). Assays such as Visium (2), Slide-seq (3) and DBiT-seq (4), use low-resolution (e.g. units larger than a cell) spatial barcoding followed by sequencing. More recent sequencing-based approaches, such as Visium HD (5) and Open-ST (6) reached subcellular resolution, which greatly increased the amount of output data. Another category of spatial transcriptomics assays are based on combinatorial barcoding, probe-hybridisation and multiple rounds of image acquisition, and include Xenium (7), CosMx (8) and MERSCOPE (9). These imaging-based spatial transcriptomics technologies have enabled the profiling of up to millions of single cells with gene panels of 200-1000 genes (7, 8), already producing data at a previously unprecedented scale. Such technologies are going to soon cover the whole transcriptome, while the area of assayed samples and the number of experimental replicates is also expected to further increase.

Multiple end-to-end and task-specific tools have been built in R and Python to analyse spatial transcriptomics data (10–12), using iterative and interactive workflows established in single-cell biology (13). Following pre-processing, specific analysis tasks include identifying spatially variable genes (using various geospatial statistical methods (12)), classifying and localising cell types or states, identifying spatial domains (14–17), and predicting ligand-receptor interactions (18–20). Notably, none of these tasks, except for direct localisation of cells, requires single-cell resolution. Given the size of high-resolution datasets, especially when multiple replicates are obtained, even quality control and exploratory analysis have become unfeasible on a laptop. Dataset size is therefore a significant bottleneck for those without a computational background and access to high-performance computing resources.

In single-cell biology, various algorithms have been developed to downsample large datasets for memory efficiency, while keeping key characteristics (21, 22). For spatial data, maintaining 2D relationships is essential, therefore completely omitting datapoints must be avoided. Instead, downsampling can be performed by reducing resolution through binning neighboring transcripts. A previous approach called SEraster binned transcripts into squares, which speeds up computation of spatially variable genes, but loses cellular resolution (23). SEraster suffers from multiple weaknesses: firstly, it requires the dataset to be loaded into memory, meaning that it does not solve the bottleneck of initial dataset size. Additionally, rather than producing raw files, its output is a processed SpatialExperiment object (24). This limits users to stay within the R/Bioconductor framework, preventing access to tools developed in the Python/scanpy framework. Thirdly, binning via squares, as also done in the SEraster pre-print (23) and other studies (25), produces a sub-optimal tessellation compared to a hexagonal lattice. A hexagonal tessellation decreases edge-effects, provides a more natural visualisation, represents curved features more accurately, and has a clearly defined nearest-neighbor assignment, something that is rather ambiguous for square tessellations (26–28). Hexagonal tessellations are available in the R package Giotto Suite (29), and the most recent version of SEraster (30), however, these approaches still suffer from requiring the full dataset in memory and restricting users to the R/Bioconductor framework.

To address these shortcomings, we developed a suite of tools in Python called Pseudovisium, that replicates the output of the 10x Space Ranger pipeline for Visium data (31), except using high-resolution spatial data as input (Fig 1). This is achieved using the *pseudovisium_generate* module, that performs hexagonal binning of transcripts from high-resolution spatial data, and mimics the structure of Visium arrays (Fig 1). This in turn enables an order of magnitude more memory-efficient and faster analysis. Notably, this method is segmentation-agnostic, and therefore also captures transcripts not assigned to cells. From a Pseudovisium folder, one can load the data to any of the widely-used analysis tools (10, 11, 32) or call the *pseudovisium_qc* module to generate a .html quality-control report (Fig 1). This report contains summary statistics and visualisations on sample and probe quality, and compares transcript abundance and spatial variation between datasets. Finally, *pseudovisium_merge* can merge datasets (experimental replicates commonly) and attached tissue images into a shared coordinate system, thereby facilitating joint visualisation and side-by-side comparison of datasets throughout exploratory analysis (Fig 1). Unlike existing tools, Pseudovisium efficiently pre-processes the data to eliminate memory bottlenecks, and generates raw data compatible with R and Python frameworks.

**Figure 1.**
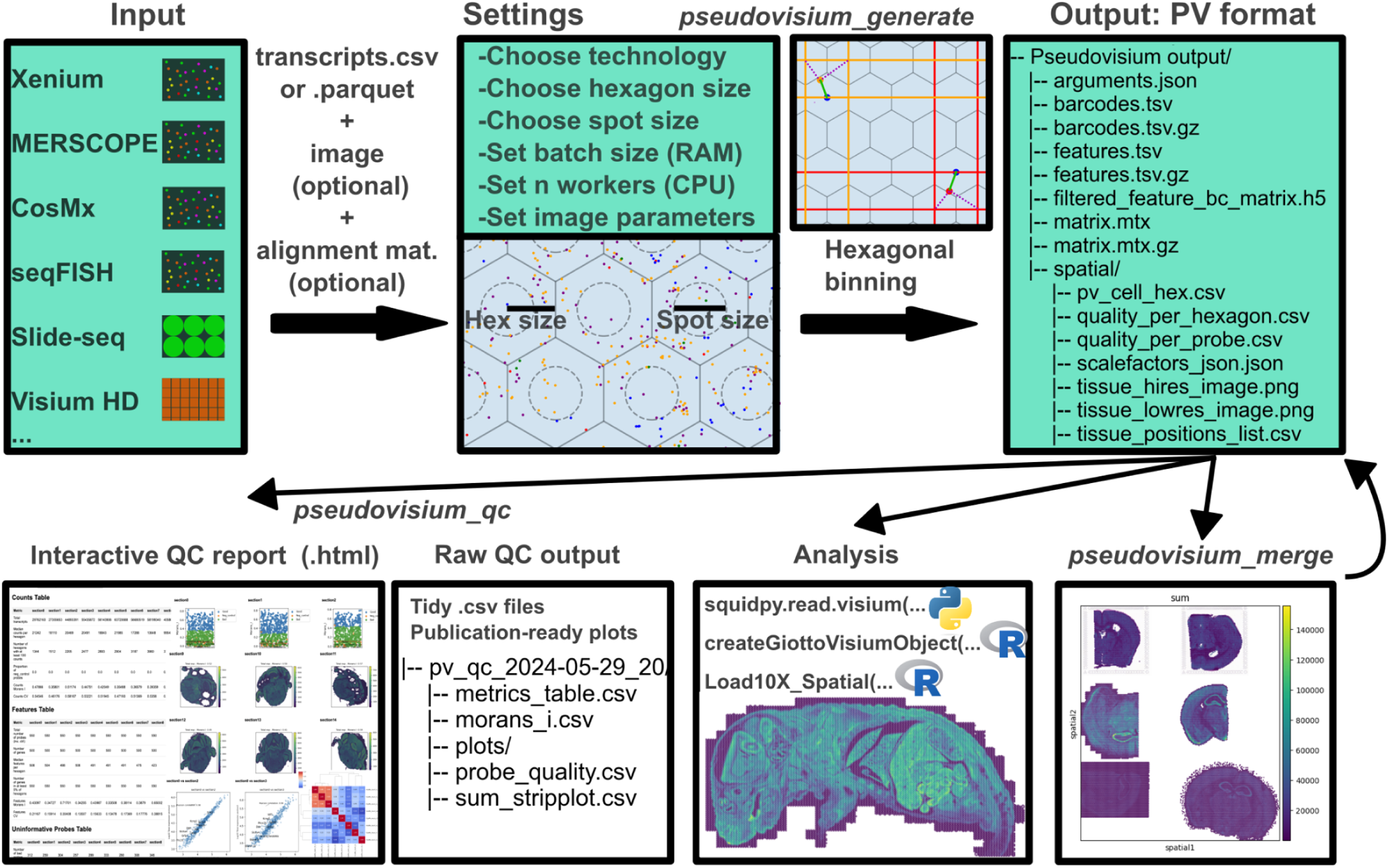
: Scheme showing the Pseudovisium (PV) workflow. *Pseudovisium_generate* takes a transcripts.csv file (or equivalent, see Methods), optionally a tissue image and corresponding alignment matrix, as well as multiple parameters to perform hexagonal-binning and generate a PV-formatted folder. Two examples (orange and red) of how transcripts are binned into hexagons are shown. In each case, a bounding square is found, and of the three possible hexagon centres, the closest one is assigned (marked by green line). The structure of the PV output folder resembles that of the 10x Space Ranger output (31). From the PV folder, the user can generate a QC report using *pseudovisium_qc*, or read in the data to established analysis tools such as squidpy (11), Giotto (10) or Seurat (32). Alternatively, multiple PV folders can be merged to a single dataset using *pseudovisium_merge*.

Here, using 47 publicly available datasets, we first demonstrate how Pseudovisium enhances the speed and memory efficiency of downstream exploratory analysis, with minimal trade-offs in results compared to using high-resolution data. Additionally, we demonstrate that applying Pseudovisium to a high-resolution input dataset can effectively mimic a Visium experiment. Furthermore, we highlight the utility of rapid quality control on three large-scale spatial datasets: a CosMx non-small cell lung cancer dataset (8), a Xenium pulmonary lung disease dataset (33), and a mouse whole-brain MERSCOPE dataset (34). We then convert 52 publicly available high-resolution datasets to Pseudovisium format and compare 6 spatial transcriptomics technologies and contrast their abilities on transcript capture. We release all generated Pseudovisium folders as a community resource (Supplementary file 1). Finally, we demonstrate how Pseudovisium allows the merging of datasets from different technologies for joint analysis of brain regions.

## Results

### Pseudovisium increases speed and memory efficiency

We first tested Pseudovisium (PV) with 100 µm diameter hexagons on a CosMx mouse brain spatial transcriptomics experiment (35), containing 48,011 cells and 116 million transcripts. The hexagonal binning resulted in a 9 times smaller AnnData object (36), with a 25 times smaller dense matrix. To evaluate the conservation of biological signals, we compared the PV data with the high-resolution ground truth in terms of clustering, spatially variable genes, spatially enriched gene sets, gene-gene correlations, and marker-marker co-occurrence (Methods). Our analysis revealed moderate agreement in clustering (Adjusted Rand Index: 0.44; Fig 2A), and strong correlation in the ranks of spatially variable genes (Geary’s C - Spearman r: 0.85; Moran’s I - Spearman r: 0.87; Fig 2B) and gene sets (Moran’s I - Spearman r: 0.81; Fig 2C). Among the genes and gene sets with highest spatial autocorrelation, we highlight *Mbp* (Fig 2B) and the long-term potentiation gene set from Reactome (37) (Fig 2C). These results demonstrate that PV effectively preserves critical spatial information despite reducing dataset size. While consistent in ranks, both Moran’s I and Geary’s C statistics were inflated in the PV data compared to the high-resolution version (Fig 2B). Similarly, the directions and ranks of gene-gene correlations were conserved in the PV data, but their magnitude was inflated (Fig 2D). Additionally, binning of the data did not result in mixing of marker genes, as highlighted by the similar frequency of marker overlap between the PV and high-resolution data (Fig 2E). This indicates that PV maintains the integrity of marker gene expression patterns, which is crucial for accurate biological interpretation.

**Figure 2.**
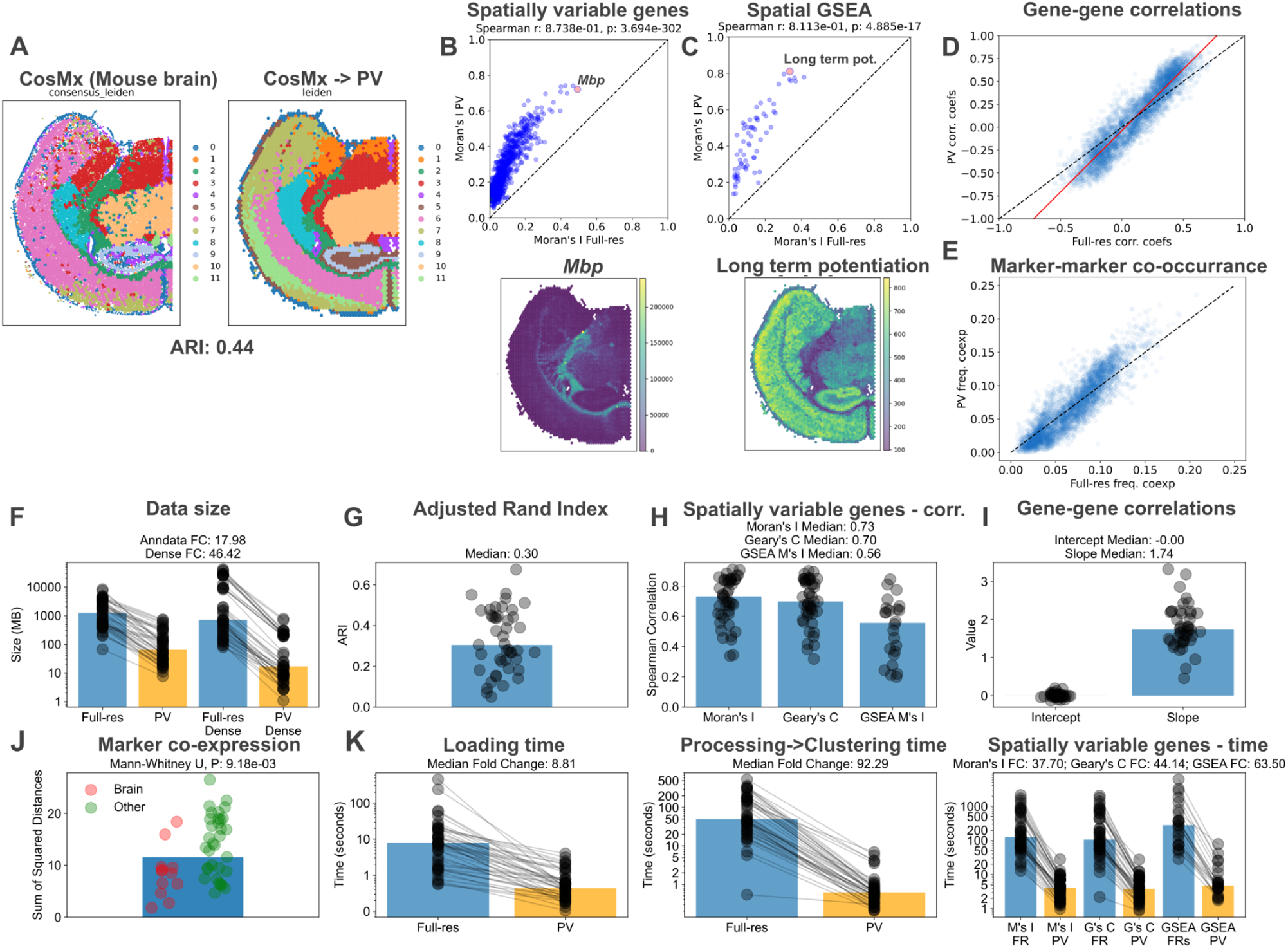
: Analysis of 47 datasets shows that *pseudovisium_generate* module successfully compresses data 10 to 50-fold and speeds up analysis by 10 to 100-fold. A: 2D spatial plot of mouse brain cells assayed by CosMx and the generated Pseudovisium hexagons colored by their respective Leiden cluster assignments. B: (Top) Scatterplot of spatially variable genes in the Pseudovisium data compared to the original high-resolution dataset. The dotted line marks the diagonal. (Bottom) 2D spatial plot of the top spatially variable gene *Mbp*. C: (Top) Scatterplot of spatially enriched gene sets in the Pseudovisium data compared to the original high-resolution dataset. The dotted line marks the diagonal. (Bottom) 2D spatial plot of the top spatially enriched gene set “Reactome Long-term potentiation”. D: Scatterplot showing the gene-gene Spearman correlation coefficients in the Pseudovisium data compared to the original high-resolution dataset. The black dotted line marks the diagonal, while the red line shows the fitted linear model with a slope > 1. E: Scatterplot showing the frequency of genes being highly expressed markers (above 75th percentile) in the same hexagon (Pseudovisium) versus in the same cell (high-resolution original data). The black dotted line marks the diagonal. F: Stripplot-barplot showing the data size (AnnData object, and dense matrix) of various high-resolution datasets and their Pseudovisium counterparts on a log scale. G: Stripplot-barplot showing ARI between the clustering of various high-resolution datasets and their Pseudovisium counterparts. H: Stripplot-barplot showing the Spearman correlation coefficients in spatially variable genes (Moran’s I and Geary’s C in squidpy (11)) and spatially enriched gene sets (spatialAUC - Methods) between various high-resolution datasets and their Pseudovisium counterparts. I: Stripplot-barplot showing the intercept and slope values of linear models derived from the relationship of gene-gene correlations between various high-resolution datasets and their Pseudovisium counterparts. The intercept near zero, and slope > 1 argues for consistently inflated gene-gene correlation values. J: Stripplot-barplot showing the sum of squared distances from the relationship of marker-marker coexpression between various high-resolution datasets and their Pseudovisium counterparts. The red color marks brain samples, and green color marks all other samples. K: Panel of plots showing that Pseudovisium increases the speed of analysis with comparable results to high-resolution datasets. (Left) Stripplot-barplot showing the loading time into an AnnData object for various high-resolution datasets and their Pseudovisium counterparts on a log scale. (Middle) Stripplot-barplot showing the time taken to process and cluster (Methods) various high-resolution datasets and their Pseudovisium counterparts on a log scale. (Right) Stripplot-barplot showing the time taken to calculate spatially variable genes (Moran’s I and Geary’s C in squidpy (11)) and spatially enriched gene sets (spatialAUC - Methods) on various high-resolution datasets and their Pseudovisium counterparts on a log scale. ARI: Adjusted Rand Index, FC: Fold-change, PV: Pseudovisium data, FR: Full-resolution data.

Loading in the data was 11 times faster, processing and clustering of the data were 100 times faster with PV, compared to the high-resolution data. Finally, computation of spatially variable genes was 15 times faster for both Moran’s I and Geary’s C respectively, and spatial gene set enrichment analysis was 13 times faster.

To generalise these findings we analysed further 46 publicly available datasets both from commercial and academic sources (Methods), including 5 technologies: 10x Xenium, 10x Visium HD, Nanostring CosMx, Vizgen MERSCOPE, Slide-seq (Curio Seeker) (Supplementary table 1; PV folders for all datasets available in Supplementary file 1). Hexagonal-binning resulted in a median 18-fold reduction in the size of the AnnData object, and a median 46-fold reduction in dense matrix size (Fig 2F). Following processing, we found a modest median ARI of 0.3, with 17 samples having an ARI above 0.4 (Fig 2G). For comparison, ARI values of 0.3-0.6 are considered state-of-the-art in spatial domain detection, when comparing against manual histology annotations (38, 39). Spatially variable genes showed high similarity between the PV and high-resolution datasets, with a median Spearman correlation coefficient of 0.73 and 0.7 for Moran’s I and Geary’s C statistics respectively (Fig 2H). For enriched gene sets, this value was slightly lower with a median Spearman correlation coefficient of 0.56 (Fig 2H). Gene-gene correlation values were generally inflated in the PV datasets compared to the high-resolution versions, illustrated by the slope values well above 1 (median 1.74; Fig 2I). To systematically compare marker-marker co-occurrence, we measured the sum of squared deviations from the values expected for each pair of markers based on the high-resolution data. Interestingly brain samples exhibited minimal mixing of markers, compared to other tissues with higher marker overlaps (Mann-Whitney U, P=9E-3; Fig 2J).

Throughout the analysis, we observed significant improvements in processing times: a median 9-fold speed-up in data loading, a 92-fold speed-up in processing and clustering, and a median 38-fold and 44-fold speed-up on spatially variable gene calculation using Moran’s I and Geary’s C (Fig 2K). Furthermore, spatial gene set enrichment analysis was also faster by a median 64-fold (Fig 2K).

Overall our results demonstrate that Pseudovisium achieves more than an order of magnitude increase in speed and memory efficiency, while preserving essential structures in clustering, and showing high concordance in spatially variable genes, gene-gene relationships and spatial enrichment of gene sets. Additionally, the observed pattern of inflated Moran’s I and Geary’s C values, as well as gene-gene correlations, may reflect a general property of low-resolution spatial data (e.g. Visium), which has not been previously explored to the best of our knowledge.

### Pseudovisium accurately simulates Visium data to inform experimental design

While Visium arrays follow a hexagonal arrangement (with hexagonal diameter of 100 µm), there is an incomplete tessellation with each spot only covering a circle with 55-65 µm diameter (31, 40). To accurately recapitulate this, Pseudovisium includes the option for users to choose a spot diameter. Previous attempts to recapitulate Visium data for benchmarking various algorithms (25, 27) and comparison of technologies (41) have been inconsistent in their approaches, given the lack of tools designed for this task.

To demonstrate that Pseudovisium can accurately simulate Visium, we turned to published datasets where consecutive tissue slices have been assayed with Visium and a high-resolution technology (Xenium (7) or CosMx (42)). By converting the high-resolution data to PV format, we could directly compare it to the actual Visium data. First, we analysed the Xenium-derived dataset (7), and we found a similar spatial distribution of clusters compared to the Visium data (Fig 3A). We then observed that PV spots contained more counts than Visium spots, reflecting the superior transcript capture of the Xenium platform (Fig 3B). Accordingly, raw expression of transcripts per spot was higher in the PV data, but it was strongly correlated with the Visium data (Fig 3C), suggesting that Pseudovisium effectively recapitulates quantitative gene expression levels. Furthermore, we observed a high correlation in Moran’s I values for normalised gene counts between the two datasets (Fig 3D), further validating the accuracy of spatial gene expression patterns preserved by Pseudovisium. Moreover, spatial gene set enrichment analysis, gene-gene correlations and marker-marker segregation also appeared strongly conserved between the PV and Visium datasets (Fig 3E).

**Figure 3.**
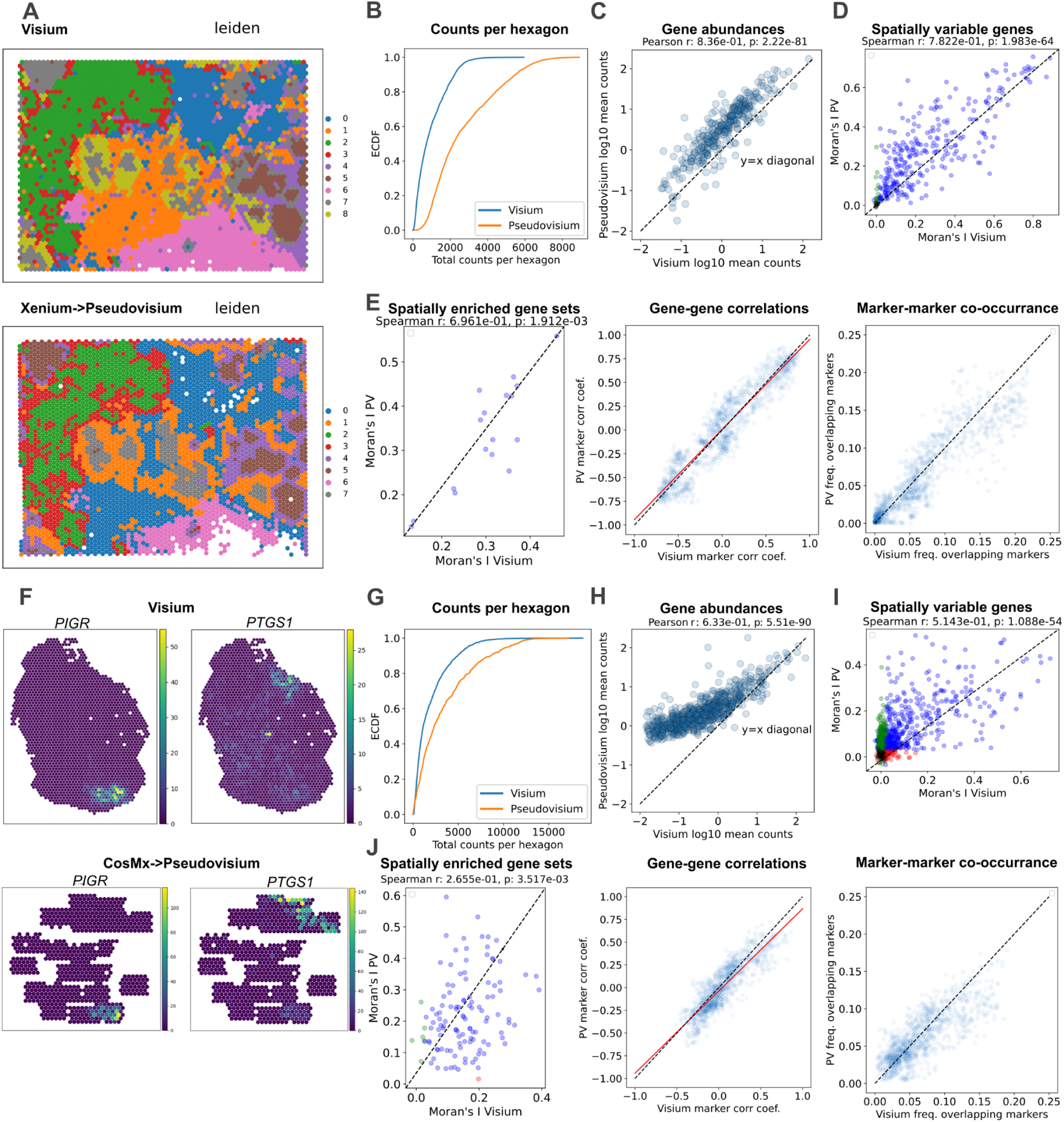
: The *pseudovisium_generate* module can accurately simulate Visium experiments when a spot diameter is included. A: 2D spatial plots colored by their separately assigned Leiden clusters. B: Empirical cumulative distribution function (ECDF) plot showing the distribution of counts per hexagon/spot in the Pseudovisium versus real Visium data. That the Pseudovisium data has higher counts shows that the underlying high-resolution technology, Xenium, has better transcript capture. C: Scatterplot showing the gene abundances in the Pseudovisium and Visium datasets. The dotted black line marks the diagonal. D: Scatterplot showing the spatial variability (Moran’s I) of genes in the Pseudovisium and Visium datasets. E: (Left) Scatterplot showing the spatial enrichment (Moran’s I) of gene sets in the Pseudovisium and Visium datasets. (Middle) Scatterplot showing the gene-gene Spearman correlation coefficients in the Pseudovisium data compared to the real Visium dataset. The black dotted line marks the diagonal, while the red line shows the fitted linear model with slope≈1. (Right) Scatterplot showing the frequency of genes being highly expressed markers (above 75th percentile) in the same hexagon (Pseudovisium) versus in the same spot (Visium). The black dotted line marks the diagonal. F: 2D spatial plots colored by (Left) *PIGR* expression or (Right) *PTGS1* expression in the (Top) Visium versus (Bottom) Pseudovisium data. G: Empirical cumulative distribution function (ECDF) plot showing the distribution of counts per hexagon/spot in the Pseudovisium versus real Visium data. That the Pseudovisium data has higher counts shows that the underlying high-resolution technology, CosMx, has captures more transcripts. H: Scatterplot showing the gene abundances in the Pseudovisium and Visium datasets. The dotted black line marks the diagonal. The “banana-shaped” curve argues that lowly-expressed transcripts have inflated counts in the CosMx data. I: Scatterplot showing the spatial variability (Moran’s I) of genes in the Pseudovisium and Visium datasets. J: (Left) Scatterplot showing the spatial enrichment (Moran’s I) of gene sets in the Pseudovisium and Visium datasets. (Middle) Scatterplot showing the gene-gene Spearman correlation coefficients in the Pseudovisium data compared to the real Visium dataset. The black dotted line marks the diagonal, while the red line shows the fitted linear model with slope≈1. (Right) Scatterplot showing the frequency of genes being highly expressed markers (above 75th percentile) in the same hexagon (Pseudovisium) versus in the same spot (Visium). The black dotted line marks the diagonal.

Next, we repeated the analysis for the CosMx-derived dataset (42). Given the disjoint spatial structure and small covered area in the CosMx dataset, we compared gene signatures identified in the original publication, instead of clusters. We found that patterns of two marker genes discussed in the original publication (42), *PTGS1* and *PIGR*, remain identifiable following PV conversion (Fig 3F). Again, the PV dataset had higher counts per spots than the real Visium data, suggesting that CosMx also captures more transcripts (Fig 3G). However we noticed that lowly expressed transcripts in Visium showed a plateau in PV data (Fig 3H). This “banana-shaped” curve is likely due to the inaccurate identification of lowly expressed genes in the CosMx technology, which has been previously reported (43–46). Despite the lower accuracy of CosMx and its patchy spatial coverage, spatially variable genes (Fig 3I) as well as enriched gene sets, gene-gene correlations, and marker-marker segration remained somewhat conserved between the PV and Visium datasets (Fig 3J).

Lastly, given thst high-resolution ground-truth data is often missing, it can only be speculated that Visium spots usually represent 1-10 cells (47). Using the PV data, we found that a median of 19 cells contributed to the expression of a spot in both datasets. Evaluating the number of cells per spot in a pilot dataset will advise researchers on choosing the appropriate spatial platform for future experiments.

We, therefore, argue that Pseudovisium is a suitable method to mimic results of the 10x Visium technology, with use cases including algorithm benchmarking and cross-technology comparisons. Additionally, following a pilot high-resolution spatial experiment, researchers can compare their results with the PV version to decide whether or not their biological insights require high-resolution data for further experimental replicates.

### Pseudovisium format facilitates rapid quality control of large-scale CosMx, Xenium, and MERSCOPE datasets

The lightweight format of Pseudovisium is ideal for rapid exploratory data analysis, including quality control. Spatial transcriptomics experiments often include multiple replicates, from different individuals and conditions, making it desirable to perform shared QC across all replicates and compare them side-by-side. Such QC can identify low-quality samples or tissue regions, as well as faulty probes. To demonstrate this capability, we accessed high-resolution CosMx (8), Xenium (33), and MERSCOPE (34) datasets containing 8, 28, and 60 replicates respectively, to perform QC. Using the *pseudovisium_qc* module, we then generated a shared QC report for each of these datasets using a single command (the full reports are accessible as Supplementary file 2, 3 and 4).

To compare experimental replicates, the exploratory QC reports include dendrogram-heatmaps, which highlight replicates that correlate either in transcript abundance or spatial variability of transcripts (Moran’s I scores). First we analysed the CosMx non-small cell lung cancer dataset with 8 replicates, comprising approximately ∼0.8 million cells and ∼260 million transcripts (8). Gene abundance analysis showed that a group of 3 technical replicates and another group of 2 technical replicates were indeed similar, as one would expect (Fig 4A). However, when examining spatial variability of transcripts, only the 3 technical replicates remained similar, while the other 2 technical replicates appeared less correlated (Fig 4B). The QC report also includes key metrics, such as median transcript counts per hexagon, median cell density per hexagon, the number of uninformative probes (similar in distribution to negative control probes), as well as the median number of unassigned (to cells) transcripts per hexagon. These metrics showed that large variability exists between replicates, often differing by a factor of 2 in these metrics (Fig 4C).

**Figure 4.**
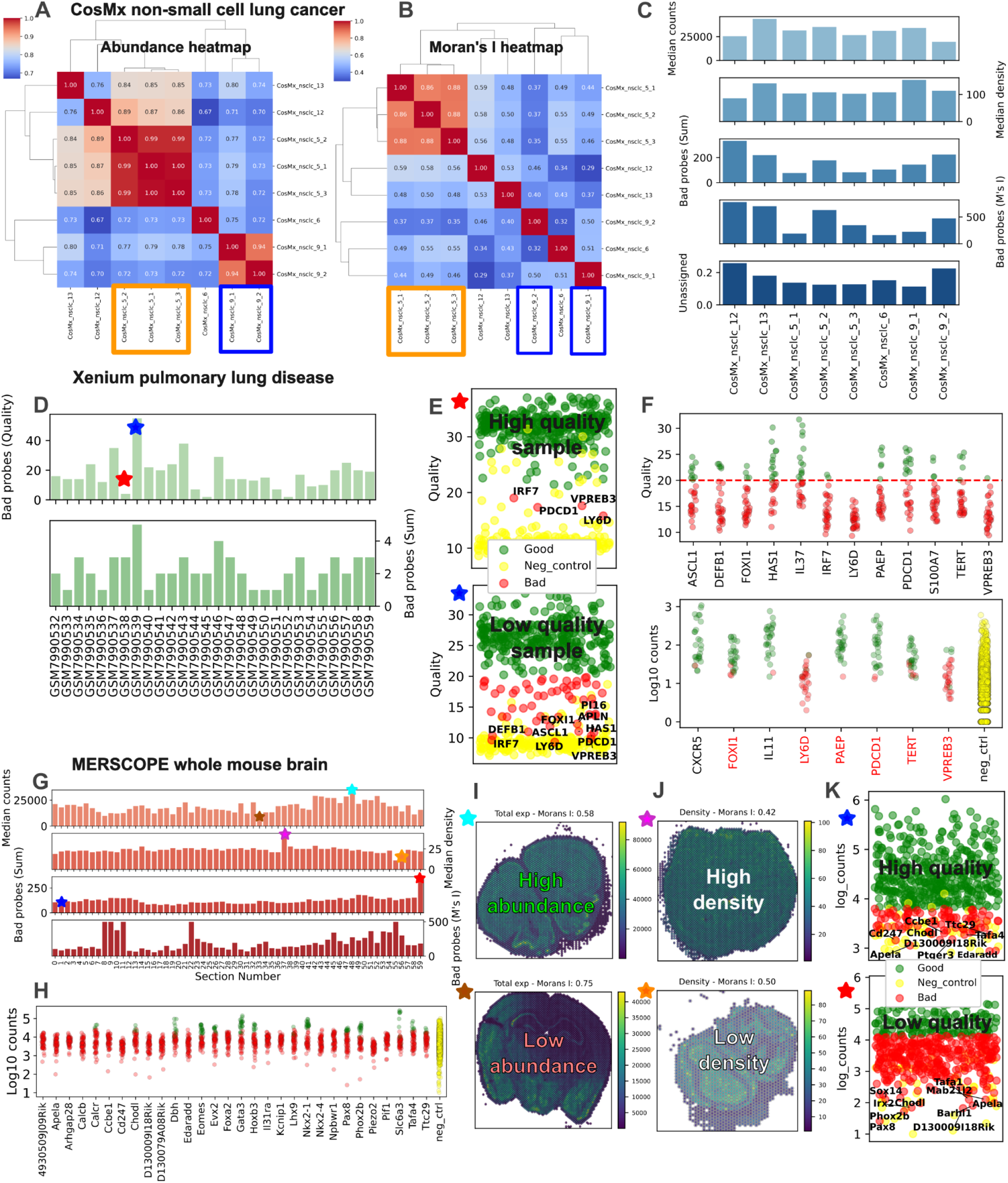
: The *pseudovisium_qc* module enables qc of three large datasets: CosMx non-small cell lung cancer (8 replicates), Xenium pulmonary lung disease (28 replicates), and MERSCOPE whole mouse brain (60 replicates) datasets. A: Dendrogram-heatmap of correlations in transcript abundance between CosMx experimental replicates. Two groups of technical replicates are highlighted with blue and orange rectangles. B: Dendrogram-heatmap of spatial autocorrelation of transcripts between CosMx experimental replicates. Two groups of technical replicates are highlighted with blue and orange rectangles. C: Barplots of summary statistics between different CosMx experimental replicates, from top to bottom: median counts, median cell density, number of bad probes based on abundance, number of bad probes based on spatial variability, percentage of unassigned (to cells) transcripts. D: Barplots of summary statistics between different Xenium experimental replicates, from top to bottom: number of bad probes based on quality score, number of bad probes based on abundance. E: Two stripplots with log10 counts on the y-axis for each assayed probe for a better and worse quality experimental replicate. Negative control probes are colored yellow, probes that are not significantly outside of the distribution of negative control probes are colored with red, while probes that are significantly more abundant than negative control probes are colored with green. F: Stripplots of Xenium quality measurements and log10 counts for the 12 consistently lowest quality probes and the top 8 lowest abundance probes. Dotted red line marks the quality cut-off of 20 used by 10x Genomics. Yellow values mark the abundance of all negative control probes across all replicates. Genes highlighted in red are shared between the two plots. G: Barplots of summary statistics between different MERSCOPE experimental replicates, from top to bottom: median counts, median cell density, number of bad probes based on abundance, number of bad probes based on spatial variability. H: Stripplots of the 30 consistently lowest expression genes and all negative control probes across all replicates. I: 2D spatial plots showing total counts per hexagon for high abundance and low abundance experimental replicate. J: 2D spatial plots showing cell density per hexagon for high density and low density experimental replicate. K: Two stripplots with log10 counts on the y-axis for each assayed set of probes for a better and worse quality experimental replicate. Same color scheme used as on Fig E above.

Next, we analysed a larger dataset of 28 replicates on pulmonary lung disease collected using the Xenium platform, totalling approximately 1.3 million cells and 210 million transcripts (33). Here we used the quality scores assigned to each transcript by the Xenium platform, which revealed a number of transcripts/probes that fall under the official quality threshold of 20 recommended by 10x Genomics, and that this greatly varies from sample to sample (Fig 4D). We highlight two samples, one where most probes are above the quality threshold of 20, and another sample where many of the probes fall below this threshold (Fig 4E). Using these quality scores, we identified 12 probes consistently below the quality threshold across replicates. In addition, further probes were flagged based on their transcript abundances compared to negative control probes (Fig 4F). We thereby present a simple approach to systematically flag bad probes in large-scale experiments.

Finally, we analysed an even larger MERSCOPE experiment, with 60 replicates spanning the entire mouse brain, totalling approximately 5 million cells and 4.6 billion transcripts (34). Replicates exhibited variability in median counts and cell density per hexagon, as well as the number of uninformative probes (Fig 4G). We identified 117 probes within the distribution of negative control probes in at least 30 experimental replicates, and we show the top 30 lowest abundance ones (Fig 4H). To illustrate the summary statistics above, we compared a pair of samples with high and low (likely damaged sample) transcript abundance (Fig 4I), a pair with high and low cell density (Fig 4J), and a pair with many versus few high-quality transcripts (Fig 4K).

Overall we found that the *pseudovisium_qc* module creates informative reports that can flag potentially low-quality samples and probes. Furthermore, it generates valuable metadata for fine-tuning experimental protocols and for modelling sample-specific differences.

### The Pseudovisium format facilitates cross-technology comparisons

With the commercialisation of multiple spatial transcriptomics platforms, various studies have aimed to compare different technologies (44–46, 48, 49). Such comparisons are often limited by technological differences, including the different formats produced by sequencing versus imaging-based platforms. Importantly, platforms also differ in whether they allocate transcripts to cells, subcellular bins or large spots, prohibiting direct comparisons. Additionally, even within imaging-based platforms, direct comparisons remain challenging due to significant differences in segmentation approaches (44–46, 48). To clearly understand the differences in the number of captured transcripts, their spatial patterns, and how these relate to negative control probes, one can compare all technologies starting by converting data to a Pseudovisium format.

To demonstrate this, we compared the same 47 datasets from 5 technologies as before (Fig 2), along with additional datasets, including ones from the 10x Visium platform, making a total of 59 datasets with approximately 257.000 hexagons and 6.5 billion transcripts (Supplementary table 1; PV folders for all datasets are available in Supplementary file 1; results in Supplementary file 5).

Importantly we did not control for sequencing depth (49) (not applicable to imaging-based methods) or sample thickness, and assumed each experiment to be the optimised output of each platform. Additional limitations include pooling probe-based and polyA-based Visium experiments together, and that datasets were not acquired from paired sections, and often are from different organs. As such, we focused on a qualitative assessment of each platform, particularly to demonstrate the utility of Pseudovisium in comparing datasets acquired by different technologies.

Overall, we found that Visium HD captures the most transcripts per hexagon, followed by CosMx, Visium and MERSCOPE, with Slide-seq and Xenium capturing the fewest (Fig 5A). Note that this analysis ignores the difference in number of genes detected (often constrained by target gene panel). The number of detected genes per hexagon was highest in Visium HD and Visium, followed by Slide-seq and CosMx, with MERSCOPE and Xenium capturing the fewest (Fig 5B). When accounting for the total number of detected genes, MERSCOPE and CosMx had the highest counts per hexagon per gene, followed by Xenium, with sequencing-based technologies, Visium HD, Visium and Slide-seq capturing the least (Fig 5A). These findings are consistent with our previous result on tissue matched Xenium-Visium (7) and CosMx-Visium experiments (42), where higher counts were observed for Xenium and CosMx compared to Visium (Fig 3B, G).

**Figure 5.**
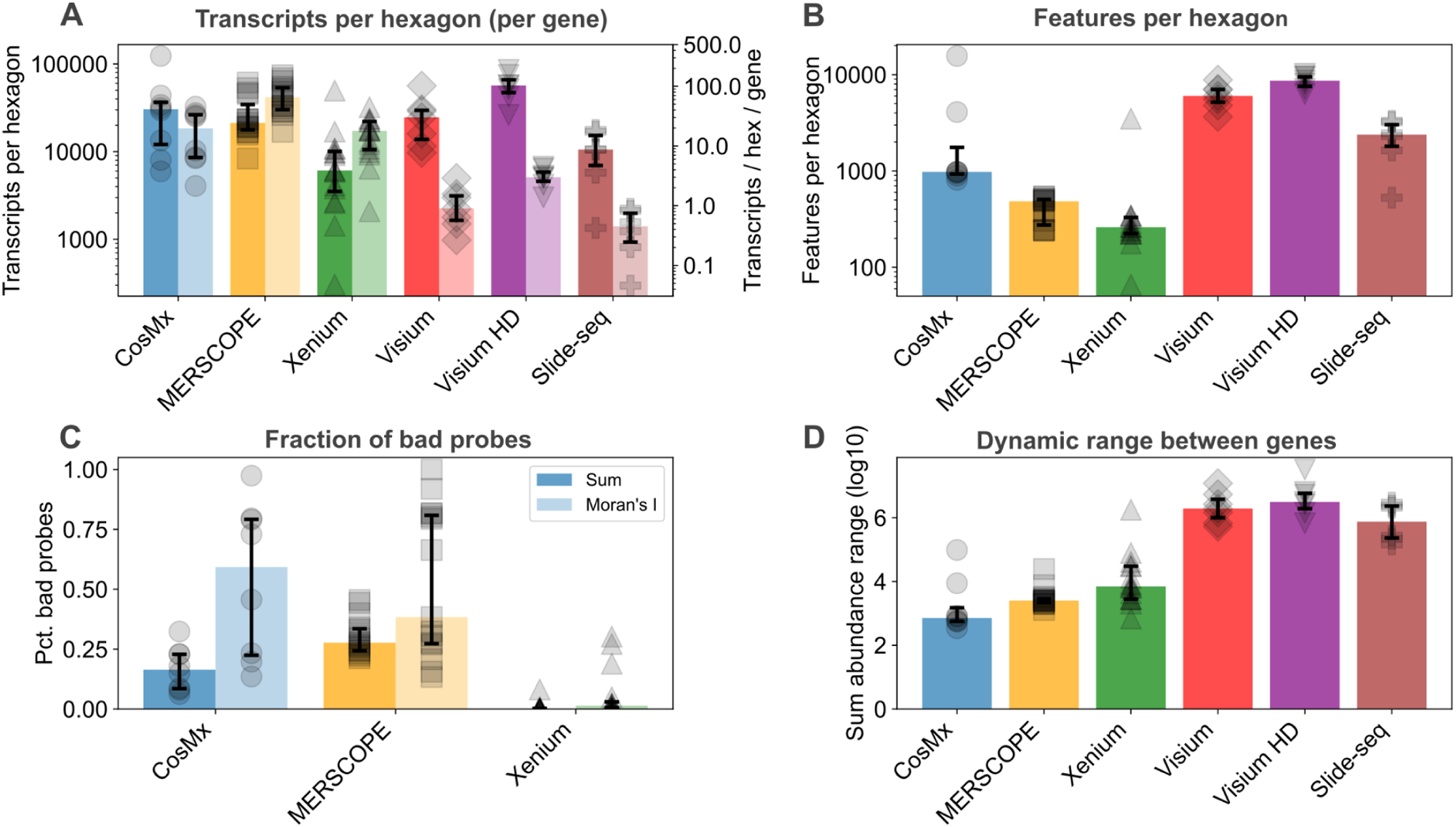
: The *pseudovisium_qc* module enables comparison of metrics across technologies. A: Double stripplot-barplot of median transcripts per hexagon and median transcripts per hexagon per gene values in each dataset for a given spatial transcriptomics platform. Error bars represent the interquartile range. B: Stripplot-barplot of features per hexagon values in each dataset for a given spatial transcriptomics platform. Error bars represent the interquartile range. C: Double stripplot-barplot of the percentage of bad probes based on abundance or spatial variability in each dataset for a given spatial transcriptomics platform. Error bars represent the interquartile range. D: Stripplot-barplot of log10 total abundance dynamic range values in each dataset for a given spatial transcriptomics platform. Error bars represent the interquartile range.

Regarding the number of uninformative probes, only technologies with negative control probes were taken forward. Based on transcript abundance, Xenium showed remarkable specificity with almost all datasets showing a near 0% of uninformative probes (Fig 5C). Conversely, most MERSCOPE datasets had 25-30% of their probes being similarly abundant as negative control probes, and CosMx had about 15-20% uninformative probes (Fig 5C). The results were more striking when uninformative probes were called based on the Moran’s I statistic, often suggesting that the majority of probes fall within the distribution of negative control probes for CosMx and MERSCOPE (Fig 5C). These findings raise questions about the reliability of negative control probes in these technologies, and we do not necessarily suggest that all gene-targeting probes are truly uninformative. Notably, Xenium consistently showed a clear separation in spatial autocorrelation between gene-targeting probes and negative controls (Fig 5C).

Finally, we assessed the dynamic range of counts by comparing the highest and lowest expressing genes in the datasets. Sequencing-based technologies generally exhibited a larger dynamic range, covering a million-fold difference in gene expression between genes (Fig 5D). Among the imaging-based spatial technologies, Xenium had the highest such dynamic range, sometimes reaching 10000-fold differences in transcripts, followed by MERSCOPE and finally CosMx (Fig 5D). This is in agreement with the inaccurate detection of lowly-expressed transcripts in the CosMx platform, highlighted on Fig 3H.

### The Pseudovisium format allows merging of datasets from different platforms

Most spatial transcriptomics workflows are designed for the analysis of a single dataset, typically starting from a single folder of input files. However, many experiments contain multiple replicates, necessitating their merging for joint analysis. This is particularly challenging for spatial data compared to single-cell datasets, as spatial and imaging information also has to be merged alongside cell-by-gene matrices.

Additionally, various spatial technologies have produced data for the same type of tissue sample (e.g. brain, breast cancer), but these have not been jointly analysed due to structural differences. To demonstrate such joint analysis, we used two Visium datasets (50), one Visium HD dataset (51) and one CosMx mouse brain dataset (35). We first converted the latter two to Pseudovisium format. Then, using the *pseudovisium_merge* module we merged all gene expression data into a common spatial coordinate system together with attached tissue images (Fig 6A).

**Figure 6.**
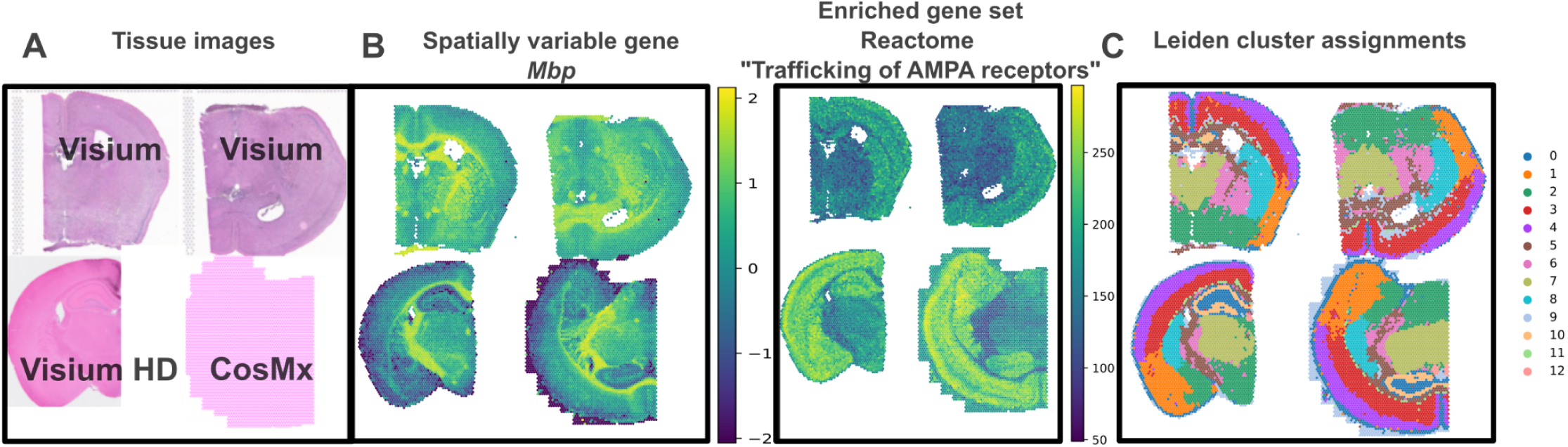
: The *pseudovisium_merge* module enables shared visualization and joint analysis of multiple spatial transcriptomics experimental replicates. A: 2D spatial plot showing the H&E images for Visium and Visium HD samples and the contours of the CosMx sample in pink. B: (Left) 2D spatial plot showing the expression patterns of one of the top spatially variable genes, *Mbp*, in the mouse brain. (Right) 2D spatial plot showing the spatial enrichment of one of the most highly enriched gene sets, “Reactome Trafficking of AMPA receptors”, in the mouse brain. C: 2D spatial plot showing Leiden cluster assignments across different samples.

Analysis of the merged object showed consistent patterns in gene expression and gene set enrichment between the different datasets (Fig 6B). We then performed batch correction using Harmony (52), and clustered using the Leiden algorithm. Remarkably, the same cell populations were revealed across the different datasets, despite having been acquired with different technologies (Fig 6C).

The capability for joint analysis will aid researchers in increasing their sample sizes by incorporating published data into their workflows, leading to more statistically robust and reproducible conclusions about the locations of cell populations and their transcriptomic signatures.

## Discussion

The recent wide-scale adoption of spatial transcriptomics necessitates novel tools to increase the findability, accessibility, interoperability and reusability of datasets. While multiple efforts have focused on increasing findability and accessibility (53–55), the other aspects have remained poorly addressed. Here, we introduce the Pseudovisium framework, which makes spatial data analysis available to wet-lab scientists by significantly decreasing the computational demand of analysis. Pseudovisium facilitates interoperability by establishing a shared format for all spatial transcriptomics data, making them compatible with a common set of analysis programs. In addition, previously written analysis scripts and software for Visium are now compatible with high-resolution datasets. The established format of Pseudovisium output (based on that of Visium), along with the decreased size of hexagonally-binned datasets, will enhance reusability of datasets, promoting both data sharing and exploration of published data.

Importantly, we find that most broad biological signatures are preserved in hexagonally-binned Pseudovisium data. Specifically, spatial signatures at the gene-level remain robustly identifiable, but accurate detection of cell type clusters depends on the degree of spatial mixing between them. Brain samples, with their distinct layered structure, preserve cell type clusters reasonably well, whereas other less organised tissues show weak conservation in cell clusters. A common problem in imaging-based spatial transcriptomics experiments is improper cell segmentation due to issues with image acquisition. After thorough benchmarking on datasets with proper segmentation, we recommend Pseudovisium as a solution to rescue such experiments, by extracting biologically meaningful conclusions through its rapid segmentation-agnostic approach. To achieve potentially more accurate results, users could decrease the hexagon size used (kept at 100 µm in this work), though this would also affect the gains in speed-up and memory-efficiency.

Our results also apply to understanding the trade-offs inherent in spatial transcriptomics platforms that are low-resolution by design (e.g. Visium, DBiT-seq). Due to a lack of sufficient high-resolution ground truth, these shortcomings of low-resolution technologies remained previously unstudied but are thoroughly explored here. Specifically, our results suggest that low-resolution technologies might be susceptible to mixing cell types and inflating spatial autocorrelation and gene-gene correlation values. While these technologies can provide a systematic first-step analysis to identify key biological players, studies in most organs will benefit from an additional cell-resolved experiment. By analogy, we suggest that most analysis could start off with rapid low-resolution step using Pseudovisium, followed-up by high-resolution analysis focusing on only the strongest hits. Future research could focus on a more thorough understanding of the trade-offs inherent in low spatial resolution, and particularly how this applies to studying various tissues.

The practical applications of Pseudovisium are extensive. Researchers can use it to integrate and analyze large-scale datasets from different platforms, enhancing the statistical robustness of their findings. Additionally, Pseudovisium’s ability to generate quality control reports will allow researchers to compare the quality of published datasets, and flag potentially corrupted samples or probes. Currently there is no such standard in sharing details on quality control, and Pseudovisium will therefore promote transparency and reproducibility of biological analysis. This quality control workflow can now be adopted by core facilities, where datasets are produced at scale.

In conclusion, Pseudovisium represents an advancement in the field of spatial transcriptomics, addressing key challenges in data analysis, quality control and cross-technology merging. By enabling more accessible analysis workflows, Pseudovisium has the potential to accelerate discoveries and foster collaborative research in this rapidly evolving field.

## Methods

### Pseudovisium package, tutorials and reproducible code

The Pseudovisium package, along with tutorials, is available on https://github.com/BKover99/Pseudovisium and downloadable from PyPI using “pip install Pseudovisium”. Additionally, Google Colab notebooks are available to reproduce all analysis, with data downloaded within them.

### Algorithm for hexagonal-binning and output of pseudovisium_generate

The main input to *pseudovisium_generate* is a flat file containing information on transcript identity, x and y coordinates, and optionally the assigned cell and quality value (e.g. for 10x Xenium). These files are always supplied as a .csv or .parquet file in the outputs of spatial platforms. The software also supports the processing of data from sequencing-based spatial transcriptomics technologies, such as Visium HD and Slide-seq (Curio Seeker). For these technologies, the input data is read from the respective file formats (e.g., Visium HD’s .h5 file or Slide-seq’s .h5ad file) and converted into a transcripts.parquet file before entering the *pseudovisium_generate* pipeline.

Given a hexagonal tessellation where one hexagon has the middle of its left edge at the origin, one can bound any point P(x,y) with a square by normalising the x coordinates with the hexagon diameter (e.g. twice the apothem), and the y coordinates with the hexagonal diameter * sqrt(3)/2. In all cases, such a square will include three hexagonal centers, in different configurations (Fig 1).

To assign transcripts to the closest hexagon, the software follows these steps:

1. Calculate the normalised coordinates (x_, y_) based on the input (x, y) coordinates and the hexagon size, and find the bounding square.
2. Determine the coordinates of the three closest hexagon centroids based on the y_ index:

○ For odd y_ indices centroids are: lower left, lower right, and upper middle.
○ For even y_ indices centroids are: lower middle, upper left, and upper right.
3. Calculate the Euclidean distances between the input point and each of the three closest centroids.
4. Select the hexagon with the minimum distance and assign the transcript to it.

The software leverages multiprocessing, which enables simultaneous processing of data batches. Importantly the user has control over batch sizes, and can adapt these for the available memory in their computing environment (Fig 1). The runtime of *pseudovisium_generate* scales linearly with the input data size. On an e2-highmem-16 virtual machine with settings batch_size=5000000, max_workers=16 we found that it took about 10 minutes to process most datasets (Supp. Fig 1).

To recapitulate the spot-array structure of real Visium experiments, the software also includes an option to only assign transcripts within a circle of set diameter around each hexagonal centroid. This is achieved by comparing the distance between the input point and the assigned hexagon centroid with the specified spot diameter. If the distance is less than half the spot diameter, the transcript is assigned to the hexagon; otherwise, it is discarded. For the entirety of the paper we used hexagons with diameter 100 µm, which captures a median of 49 cells, which for 55 µm spots translates to a median of 11 contributing cells (Supp. Fig 2).

For high-resolution sequencing-based technologies, such as the recent Visium HD (5), where transcripts are assigned to 2 µm^2^ square bins, the software has the option to smooth such values into 4 smaller 0.5 µm^2^ bins reducing edge-effects. The smoothing process is performed by distributing the count value of each transcript equally among the four neighboring bins (top-left, top-right, bottom-left, and bottom-right) within a specified smoothing distance (0.5 µm by default). This helps to mitigate the sharp transitions between adjacent hexagonal bins and provides a more continuous representation of the spatial expression patterns.

If an image file path is provided, Pseudovisium processes the image to generate high-resolution and low-resolution tissue images for visualization purposes. The image is first resized according to the specified tissue_hires_scalef parameter, which determines the scaling factor for the high-resolution image. The resized image is then transformed using an alignment matrix (affine transform matrix), which can be provided to correct for any spatial misalignment between the image and the spatial data. If the pixel_to_micron parameter is set to True, the image is rescaled based on the image_pixels_per_um parameter to convert pixel coordinates to micron coordinates. Finally, the transformed image is saved as tissue_hires_image.png and a lower resolution version (10% of the high-resolution size) is saved as tissue_lowres_image.png in the output spatial folder.

Once transcripts are assigned to hexagons, the output is organized in a folder structure that mimics the Visium datasets produced by the 10x Space Ranger pipeline (31) (Fig 1). The main output folder has the following content:

- ***filtered_feature_bc_matrix.h5***: The count matrix in HDF5 format, compatible with popular single-cell and spatial analysis tools.
- ***matrix.mtx.gz***: The count matrix in compressed Matrix Market format.
- ***features.tsv.gz***: A compressed tab-separated file containing information about the features (genes).
- ***barcodes.tsv.gz***: A compressed tab-separated file containing the hexagon barcodes.
- ***spatial/***: A subfolder containing spatial information and optional tissue images.

Within the ***spatial/*** subfolder, one can find the following files:

- ***tissue_positions_list.csv***: A comma-separated file containing the array and pixel coordinates of each hexagon in the dataset.
- ***scalefactors_json.json***: A JSON file containing information on how the hexagon coordinates scale to the original source image (if provided) and to the attached high-resolution and low-resolution images.
- ***tissue_hires_image.png*** and ***tissue_lowres_image.png***: High-resolution and low-resolution tissue images, respectively.

Additionally, the output folder includes Pseudovisium-specific files:

- ***arguments.json***: A JSON file containing all the arguments used to run the *pseudovisium_generate* command.
- ***quality_per_probe.csv*** and ***quality_per_hexagon.csv*** (optional): Files containing quality metrics for each probe and hexagon, respectively, when quality measurement available (e.g., for 10x Xenium data).
- ***pv_cell_hex.csv*** (optional): A file recording the cells (or barcodes) and their assigned counts for each hexagonal bin, generated only when cell-level information is available.

### Generation of quality control report

To assess the quality and characteristics of the Pseudovisium output, we developed a Python script that generates an interactive HTML quality control report (Fig 1). The script takes as input one or more folders containing Pseudovisium or Visium output and calculates key metrics for each dataset, including median counts and features per hexagon, proportion of negative control probes, spatial autocorrelation, and the number of probes identified as uninformative outliers. “Good” and “bad” probes are identified using a one-tailed Z-test, based on the null distribution from negative control probes, followed by Benjamini-Hochberg correction. A “good” probe is called when we reject the null hypothesis (e.g. P_adj < 0.05), otherwise the probe is flagged as “bad” and uninformative. If cell information and quality scores are available, additional metrics are calculated.

The script generates informative visualizations, such as hexagon plots displaying the spatial distribution of gene expression, unique features, and total counts. Comparative visualizations, including scatterplots and correlation heatmaps, are used to assess the correlation of probe abundance and spatial autocorrelation values (Moran’s I) between datasets. Stripplots of probes depict the distribution of quality, abundance and spatial autocorrelation, and highlight outliers.

This tool enables rapid quality assessment and comparison of Pseudovisium datasets, aiding in the identification of potential artefacts prior to downstream analysis. The HTML format facilitates collaborative sharing of the QC report as a standardized deliverable.

### Merging datasets

*Pseudovisium_merge* takes as input multiple folders containing either Pseudovisium or Visium output files, loads the data from each folder, and extracts a common scale factor (Fig 1). It concatenates the barcodes (e.g. hexagons or spots) and features (e.g. transcripts), creates a consensus index for the features across all datasets, and adjusts the count matrices and barcodes to use the consensus feature indices and a cumulative barcode indexing.

To visualize the merged data, the tool stitches the tissue images from all datasets into a single large image, arranged in a grid. The tissue positions are updated to reflect the new coordinates in the stitched image.

Finally, it saves the merged output in the above defined Pseudovisium format, including the count matrix, features, barcodes, and stitched tissue images. If cell-level information is available, the merged pv_cell_hex file is also saved.

### Datasets

Datasets were obtained from commercial and published sources and were chosen to represent a broad range of tissues and experimental sizes, both in mouse and human. Commercial sources included the 10x Genomics (https://www.10xgenomics.com/datasets), Vizgen (https://vizgen.com/data-release-program/), Nanostring (https://nanostring.com/products/cosmx-spatial-molecular-imager/ffpe-dataset/) websites. Published datasets included the Xenium pulmonary lung disease dataset from (33), the mouse whole-brain MERSCOPE dataset from (34), a high-grade glioma CosMx and Xenium dataset from (56), an ovarian cancer CosMx and Visium dataset (15), ageing brain Visium dataset from (50), a heart MERSCOPE dataset (57) a brain Slide-seq dataset (58), and an ovary Slide-seq dataset (59).

All datasets used in the paper are listed in Supplementary table 1 with download links provided, and PV folders outputted by *pseudovisium_generate* are found in Supplementary file 1, providing a rich resource for further systematic studies.

### Data processing

While we acknowledge that there is no one-size-fits-all approach that works best for all datasets, we followed a workflow similar to that commonly used in single-cell RNA-seq. First, each dataset was loaded into squidpy/scanpy (11, 60), following which sparsely expressed genes and low-quality cells were filtered out. For datasets with more than 2000 genes, we then continued with the top 2000 highly variable genes (calculated in scanpy using flavor=”seurat_v3”) to decrease the computational burden of the high-resolution datasets. This was followed by total count normalisation, applying log1p, scaling, principal component analysis, construction of KNN-graph on the first 20 PCs, and Leiden clustering. To get a comparable number of clusters between Pseudovisium and high-resolution versions of the same dataset, the resolution parameter was fixed for the high-resolution dataset, and was varied for the Pseudovisium dataset until getting a similar number of clusters.

To evaluate similarity in clustering, first the most common high-resolution cluster value was chosen for cells falling into each hexagon (“majority vote”), and was compared with the cluster assignments called for hexagons. Similarity was measured using the Adjusted Rand Index via the sklearn.metrics implementation (61). Briefly, the Adjusted Rand Index is a measure of the similarity between two sets of labels (clustering in this case), adjusted for the chance grouping of elements. A value close to 0.0 indicates random labeling, while increasing values suggest an overlap in labels, and 1.0 means identical labels.

Spatial neighbours were called using a 250 µm radius around each cell/spot/hexagon. Spatially variable genes were identified by the widely used Moran’s I and Geary’s C metrics for spatial autocorrelation, calculated using the implementation within squidpy (11). Briefly, Moran’s I and Geary’s C measure how well the expression of an index cell, spot or hexagon can be predicted by the expression of its neighbors. When spatial autocorrelation is high, Moran’s I approaches 1 and Geary’s C approaches 0.

Pairwise gene-gene correlations were measured for the top 100 most spatially variable genes, using the scipy (62) implementation of Spearman correlation. Marker-marker co-occurrence was measured on the same top 100 genes, by evaluating the frequency of two genes marking (above 75th percentile in expression) the same cell or hexagon. To systematically compare the results for marker-marker co-occurrence, we measured the sum of squared distances between the high-resolution and Pseudovisium values.

### Spatial GSEA using spatialAUC

Spatial gene-set enrichment (GSEA) was done using a bespoke approach we named spatialAUC, which operates similarly to AUCell developed for scRNA-seq data (63). Briefly, spatialAUC first retrieves gene sets from MSigDB using gseapy (64). We chose to exclude gene sets with less than 50% of its members or less than 5 genes represented in the spatial data. GSE scores were then obtained for each cell or Pseudovisium hexagon by:

- Creating an x-axis where genes are ordered by expression
- Creating a y-axis to measure the cumulative frequency of recovered genes from the target gene set.
- Calculate the area under the curve using integration by the trapezoidal rule implemented in numpy (65).

After obtaining enrichment scores, spatialAUC wraps the Moran’s I implementation in squidpy (11) to identify spatially enriched gene sets. We compared the resulting Moran’s I scores between Pseudovisium and high-resolution data.

SpatialAUC is available on https://github.com/BKover99/SpatialAUC and downloadable from PyPI using “pip install spatialAUC”.

### Merging of brain datasets

Datasets were merged using the *pseudovisium_merge* module and only shared genes were retained. Following this, spatial autocorrelation (Moran’s I) was carried out for each subset of data independently, and only those genes were retained that were in the top 500 spatially variable genes within each dataset independently. After PCs were calculated, Harmony (52) was applied for batch correction, which converged after 11 iterations. The KNN graph was constructed on the first 20 adjusted PCs, and the Leiden algorithm was used for clustering.

### Evaluating execution time and memory consumption

The time taken for each function to complete was measured using the time.time() function. The size of each dataset was evaluated using the pympler.asizeof.asizeof() function, while the memory consumption of dense matrices was obtained using the nbytes() method in numpy (65).

### Computing environment

All computation (and time measurements) was carried out on an e2-highmem-16 virtual machine from Google Cloud, with 16 cores and 128GB RAM, the latter of which was necessary for analysing the large high-resolution datasets.

## Acknowledgements

B.K. was funded by the Wellcome Trust as part of the Advanced Therapies for Regenerative Medicine Wellcome Trust PhD Programme (218461/Z/19/Z). A.V. was funded by the MRC (MR/S023747/1), BRC (BB/V006290/1), Medical Research Foundation (MRF-176-0002-RG-FLOH-C0929) and British Heart Foundation. We thank Jelmar Quist for early feedback on the project, and for reading a shortened draft of the manuscript, as well as other members of the King’s Spatial Biology facility for early discussions.

## Supplementary data

All supplementary data has been deposited to Figshare at https://figshare.com/articles/dataset/_b_Rapid_and_memory-efficient_analysis_and_quality_control_ of_large_spatial_transcriptomics_datasets_b_/26359984.

## Supplementary Figures

**Supp. Fig 1.**
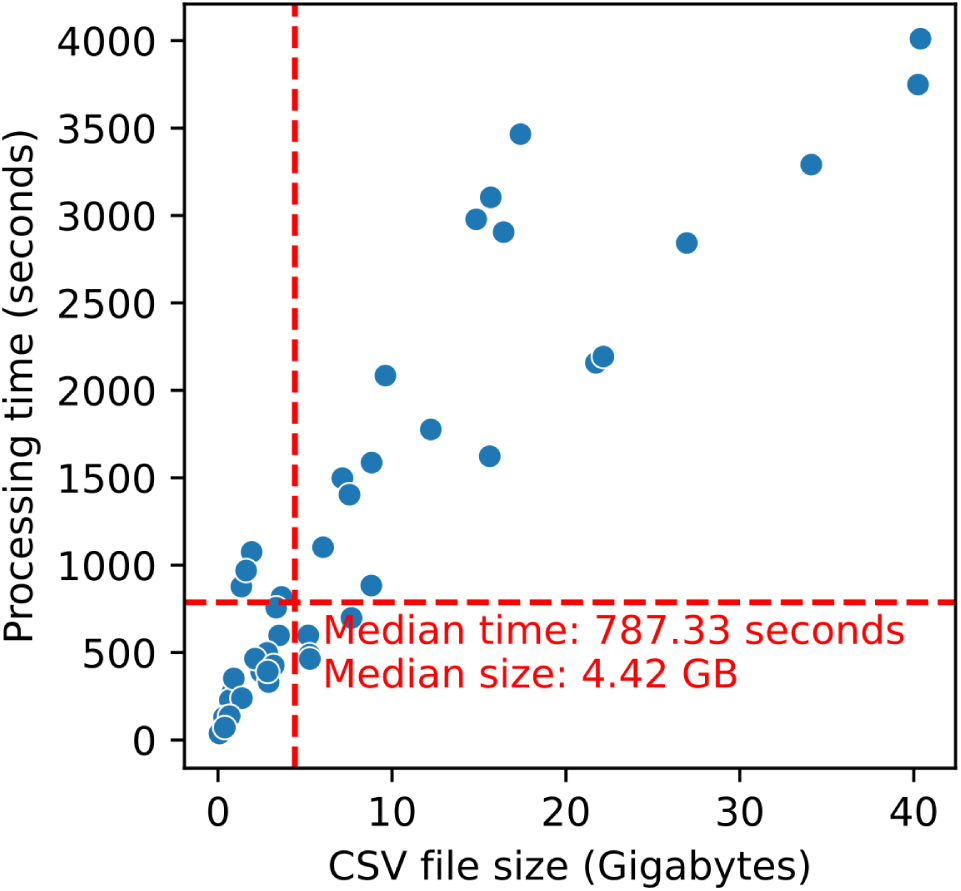
Runtime of *pseudovisium_generate* across datasets. Scatterplot showing the runtime of *pseudovisium_generate* across tested datasets and its relationship with input transcripts.csv file size. The median elapsed time and file size are highlighted in red.

**Supp. Fig 2.**
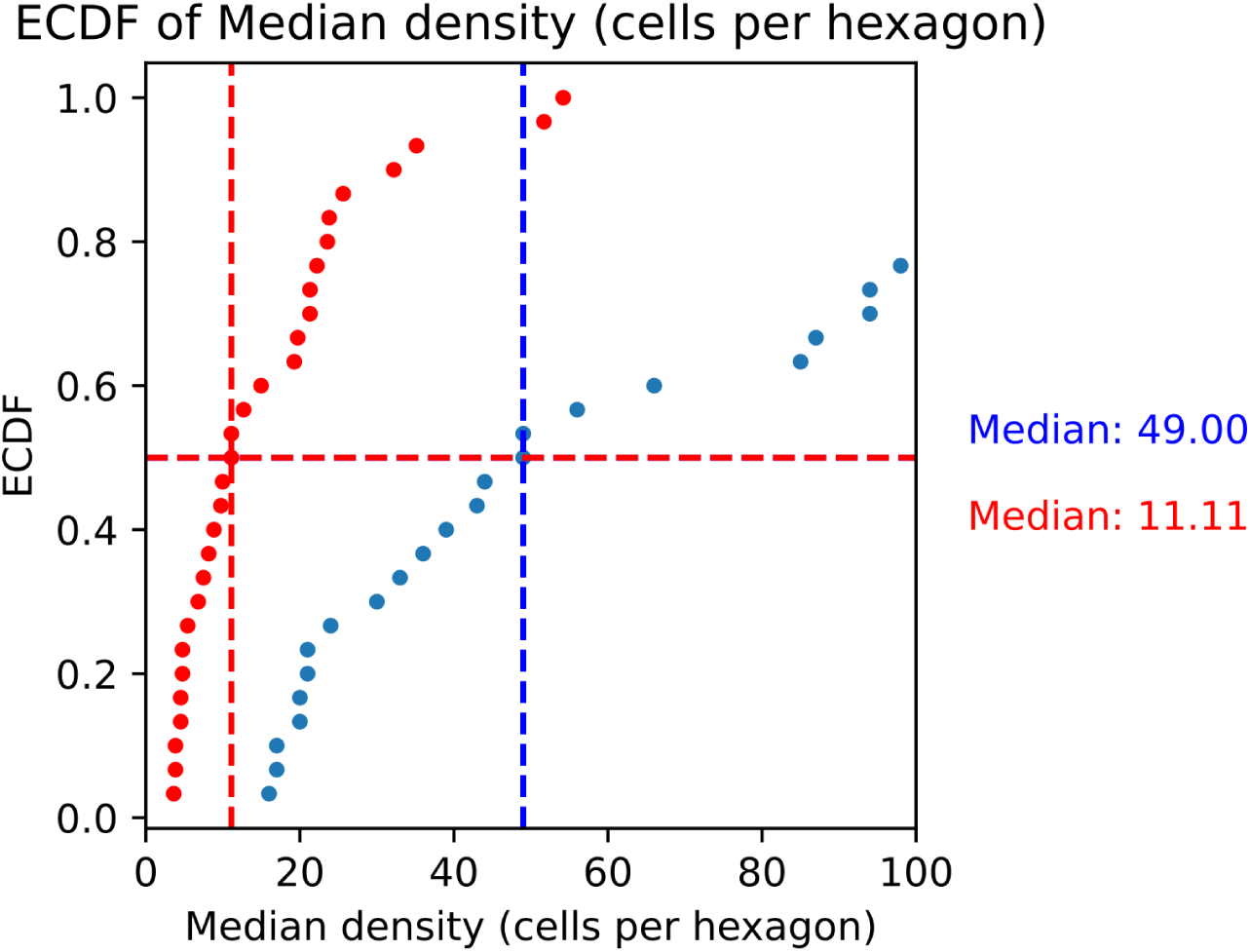
Empirical cumulative distribution function of the number of cells contributing to counts in a hexagon, and inferred for a 55 µm diameter spot (same as Visium). Each dot is a median value for a given dataset. Medians values across all datasets are highlighted in blue for hexagons and red in spots.

